# Limited directional selection but coevolutionary signals among imprinted genes in *A. lyrata*

**DOI:** 10.1101/2024.08.01.606153

**Authors:** Audrey Le Veve, Ömer İltaş, Julien Y. Dutheil, Clément Lafon Placette

## Abstract

Genomic imprinting is a form of gene regulation leading to the unequal expression of maternal and paternal alleles. The main hypothesis invoked to explain the evolution of imprinted genes is the kinship theory, which posits a conflict between parental genomes over resource allocation in progeny. According to this theory, such conflicts select for parent-of-origin–dependent expression of genes involved in resource allocation. How such conflicts translate into signatures of selection at coding or regulatory sequences remains model-dependent and is not explicitly predicted by the kinship theory. However, most studies addressing selection in imprinted genes in flowering plants, particularly those based on population-genomic or phylogenetic analyses, have focused on self-fertilizing species, where conflicts over resource allocation are predicted to be weak. Consequently, the impact of the kinship theory on the evolution of imprinted genes remains largely unexplored in systems where parental conflict is expected to be strong. Furthermore, potential coevolution between antagonistically acting imprinted genes, as proposed in extensions of parental conflict models, has not yet been tested empirically.

Using combined phylogenetic and population genomic approaches, we investigated signatures of selection on imprinted genes across the Brassicaceae family and in autogamous and allogamous populations of *Arabidopsis lyrata*, and searched for evidence of coevolution among imprinted genes. We found that endosperm-expressed genes exhibited signals of balancing selection across Brassicaceae and within allogamous populations, consistent with models of unresolved intralocus conflict. These population-level signals varied with the mating system, in line with expectations that parental conflict is reduced under self-fertilization. Moreover, phylogenetic analyses indicated signatures of purifying (negative) selection acting on imprinted genes. However, the population-level signatures of selection were independent of the mating system and showed limited concordance with kinship predictions, possibly due to stronger selection acting on expression than on coding sequences. Finally, we identified coevolution between imprinted genes, although not at specific sites, suggesting that interactions beyond protein sequence may contribute to this process.

## Introduction

The kinship theory, or the parental conflict theory, was first proposed by Haig and Westoby (1989) in the context of endosperm development and further developed in subsequent reviews in 1991. It posits that in species where the maternal parent nourishes progeny sired by multiple males, maternal fitness is maximized by distributing resources equally among offspring, whereas paternal fitness is enhanced by securing more resources for their own offspring.

While the kinship theory predicts that conflicts between maternal and paternal genomes select for expression of genes related to growth that depends on parent-of-origin, it does not explicitly address the expected patterns of allelic polymorphism at these loci (Haig and Graham 1991; Moore and Haig 1991; Wilkins and Haig 2001). Consistent with this framework, experimental crosses in *Arabidopsis thaliana* revealed dosage effects on seed size, although these patterns may reflect interactions between cytoplasmic inheritance, dosage imbalance, and imprinting (Scott et al., 1998). Balancing selection and elevated polymorphism have been predicted under alternative models of unresolved intralocus conflict acting on alleles with antagonistic fitness effects (Day and Bonduriansky 2004; Kasimatis et al., 2017). Whether similar signatures of balancing selection may arise at genes involved in resource allocation prior to, or independently of, the evolution of imprinting remains an open empirical question. Notably, signatures of balancing selection in genes expressed in the endosperm (the nourishing tissue of the embryo in the seed) have not been systematically characterised, despite their potential relevance for understanding the evolutionary consequences of parental conflict.

The kinship theory remains a leading explanation for the evolution of imprinted genes (Haig 1997; Spencer and Clark 2014; Patten et al., 2014). Imprinting is an epigenetic form of regulation in flowering plants and mammals in which allele expression depends on parental origin (Haig 1997; Wilkins 2011). By creating hemizygous expression of parental alleles, the kinship theory argues that imprinting evolves to resolve parental conflicts over resource allocation. Consequently, imprinted genes are expected to be disproportionately expressed in the endosperm in plants, the tissue where parental conflict operates (Haig 1997; Haig 2000). Thus, the proportion of Maternally Expressed Genes (MEGs) and Paternally Expressed Genes (PEGs) upregulated in the endosperm should exceed the genome-wide expectation derived from sampling all expressed genes. According to this hypothesis, many imprinted genes appear to be involved in resource accumulation in the endosperm and metabolic processes (Raissig et al., 2011). These theoretical expectations and previous empirical observations justify testing whether MEGs and PEGs are disproportionately upregulated in endosperm relative to random expectation.

Theoretical models of selection on one or two genes involved in resource allocation predict that: (1) imprinting sets distinct optimal expression levels for paternally expressed genes (PEGs) and maternally expressed genes (MEGs; Haig 1997); Specifically, PEGs are predicted to evolve higher optimal expression levels than MEGs, because paternal alleles benefit from increased resource acquisition, whereas maternal alleles tend to limit growth (Haig 2000; Patten et al., 2014). Accordingly, we expect a higher expression per active gene copy in PEGs, followed by MEGs and non-imprinted endosperm genes; (2) PEGs and MEGs are expected to experience selection favouring increased or reduced resource acquisition, respectively (Haig 2000); and (3) imprinting at some locus promoting growth may drive antagonistic coevolution with some function suppressing growth (Wilkins and Haig 2001). However, these models focus on *Insulin-like Growth Factor 2* (*IGF2*) and *Insulin-like Growth Factor 2 Receptor (IGF2R*) genes, which regulate foetal growth in mammals, without investigating the expected signatures of selection for endosperm genes. More recent theoretical work has further emphasized that selection associated with parental conflict may primarily act on loci controlling imprinting or gene regulation, rather than on the coding sequences of imprinted genes themselves, potentially weakening expected genomic signatures at these loci (Flintham et al., 2025).

Empirically, phylogenetic signals of directional selection have been reported for the *MEDEA* locus in the allogamous species *Arabidopsis lyrata*, but not in the autogamous sister species *A. thaliana* (Spillane et al., 2007; Miyake et al., 2009), and for PEGs at the population level in the autogamous Brassicaceae species *Capsella rubella* and *A. thaliana* (Hatorangan et al., 2016; Tuteja et al., 2019). However, MEGs did not show comparable signals, possibly due to technical challenges in detecting selection if selection primarily targets gene expression levels. Indeed, a central debate has emphasized that kinship theory primarily concerns selection on optimal expression levels, potentially acting on alleles related to regulatory mechanisms in parental genomes, rather than the level of polymorphism in the coding regions of imprinted genes (Patten et al., 2014). Selection acting on gene expression may still leave detectable traces in amino acid or codon usage patterns (Patten et al., 2014).

The incongruent signals detected in imprinted genes may also be due to polymorphism dynamics shaped by the mating system. The kinship theory assumes that progenies are sired by multiple males and that imprinted genes are preferentially expressed in the endosperm (Haig and Westoby 1991; Haig 2000; Brandvain and Haig 2005). The transition to autogamy may alter selection acting on genes, potentially leading to more intense directional selection or relaxed selection (Soliman and Coughlan 2024). However, the empirical studies cited above did not assess endosperm-specific expression of the imprinted genes and only focused on autogamous species. Consequently, the evolutionary signatures of selection on imprinted genes and the contribution of the kinship theory to their evolution remain unresolved in flowering plants.

Extensions of the kinship theory also predict a potential coevolution between antagonistic imprinted genes (Wilkins and Haig 2001; Willi 2013), yet no phylogenetic signal was detected for *IGF2* and *IGF2R* (McVean and Hurst 1997), and coevolution among endosperm-expressed genes remains unknown. Population genomic methods can detect coevolving sequences based on allele frequency variation; however, they may be biased by demographic or population structure effects. More integrative approaches based on phylogenetic comparisons can identify co-occurring changes in candidate coevolving genes (Dutheil and Galtier 2007).

Here, we test these predictions by combining expression, phylogenetic and population-genomic analyses to evaluate whether imprinted genes in *A. lyrata* show evolutionary signatures compatible with theoretical expectations derived from models of parental conflict. We compared imprinted genes in allogamous *A. lyrata* (Klosinska et al., 2016) with non-imprinted endosperm genes, using phylogenetic data from 23 Brassicaceae species and genomic data from nine allogamous populations and nine closely related autogamous populations of *A. lyrata* from North America (Ross-Ibarra et al., 2008). We tested for: (1) variations in expression patterns and/or codon usage; (2) reduced divergence and polymorphism, as expected under some models of directional selection, and (3) correlations between evolutionary rates or diversity metrics used as exploratory indicators of evolutionary covariance expected in case of coevolution. Our results indicate that endosperm-expressed genes display patterns compatible with unresolved intralocus conflict over resource allocation, and that the transition to autogamy promoted directional selection on these genes. Imprinted genes show evidence of ancient negative directional selection, codon usage bias, and substantial coevolution. However, the signals at the level of population are less compatible with the kinship theory, possibly due to stronger selection acting on expression than on coding sequences or by a recent relaxed selection on these genes in North American populations.

## Results

### General analytical framework

Parental conflict is expected to occur primarily in the endosperm. Using transcriptomic data from 19,425 genes across five tissues (leaf, root, pollen, pistil, and endosperm), we identified 2,732 specifically upregulated genes in the endosperm of *A. lyrata* among 31,073 genes analysed. Among these upregulated genes, 15 were Maternally Expressed Genes (MEGs) and 14 were Paternally Expressed Genes (PEGs) from the 27 MEGs and 46 PEGs previously identified in *A. lyrata* (Klosinska et al., 2016). Here and throughout the Results, we use the term ‘gene’ to refer to annotated nuclear protein-coding loci for which expression could be quantified from RNA-seq data. Between *A. lyrata* and *A. thaliana*, 40% of MEGs (6/15) and 50% of PEGs (7/14) identified in our dataset were conserved as imprinted (Table S1).

Throughout the Results section, we compare four categories of genes: MEGs and PEGs that are upregulated in the endosperm (referred to as “imprinted genes” if considered together), the 2,703 non-imprinted genes specifically upregulated in the endosperm (“endosperm genes”), and a set of 100 control genes, randomly sampled from the ubiquitously expressed genes, and serves as a baseline for genome-wide patterns of selection unrelated to parental conflict. The control panel was chosen following the strategy of sampling based on expression used in previous imprinting studies, rather than by matching genomic characteristics across groups. Differences in gene size, Coding DNA Sequence (CDS) proportion, and Guanine–Cytosine (GC) content among panels were therefore quantified and considered when interpreting downstream evolutionary analyses.

For each population genetic and phylogenetic statistic, genes with insufficient informative sites were excluded. Then, significance was evaluated using 10,000 bootstrap resampling matching the size of focal group framework, because this method is insensitive to difference in sample size. Rather than repeating the method in each subsection, we applied a single unified framework in which significance was assessed by comparing observed means to the central 95% of the empirical bootstrap distribution (i.e. 2.5^th^–97.5^th^ percentiles). This approach was used consistently across analyses of expression, codon usage, phylogenetic metrics, and population polymorphism. Although the figures share a similar visual layout for consistency, each statistic captures a distinct evolutionary dimension allowing direct comparison of patterns across gene categories while retaining analytical complementarity.

We first compared gene size, CDS content (%) and GC content (%) across groups, as these features may influence polymorphism patterns and generate bioinformatic artefacts. Control genes and MEGs were significantly smaller than PEGs and endosperm genes, and CDS content was significantly higher in endosperm and imprinted genes relative to control genes. GC content was also lower in endosperm and imprinted genes than in control genes (Table S2). These differences in gene size, CDS proportion and GC content were quantified and considered when interpreting all evolutionary analyses. Nevertheless, they were not corrected for directly, and we retained the control panel to be congruent with methods used in previous studies in the autogamous species *C. rubella* and *A. thaliana* (Hatorangan et al., 2016; Tuteja et al., 2019). In practice, all evolutionary comparisons evaluating predictions of the kinship theory were performed relative to endosperm-upregulated non-imprinted genes, which represent the expression-matched and biologically relevant background for imprinting analyses. The control panel was used only to describe genome-wide patterns of variation.

To evaluate whether these genomic differences could create artificial signals of selection, we performed several robustness checks. First, for population-genomic analyses, we verified that the filter based on Single Nucleotide Polymorphism (SNP) density did not preferentially exclude short, GC-poor, or low-diversity genes. Second, by comparing all statistics against empirical null distributions derived from endosperm-upregulated genes, we ensured that each test was evaluated against a biologically appropriate baseline sharing similar expression patterns, thereby reducing the influence of residual genomic confounders unrelated to imprinting or parental conflict. Finally, we compared different linear regression models using each statistic of interest as the response variable, and evaluated model support using the Akaike Information Criterion (AIC). The full model included gene group, gene length, CDS proportion and GC content as explanatory variables. We then evaluated a series of reduced models in which one parameter was removed at a time and retained the model with the lowest AIC value. This procedure was iterated until the best-supported model was identified. This approach allowed to determine (1) whether gene group remained a significant predictor in the best model and (2) whether genomic features accounted for the observed trends in the analysed statistics.

In previous study (Tuteja et al., 2019), validation of the kinship theory to explain imprinting was performed following another logic, based on enrichment, which evaluated not the global behaviour of imprinted genes but the proportion of them exhibiting extreme signatures compatible with parental conflict. Thus, we tested whether directional selection occurred more frequently in imprinted genes than in the empirical distribution of endosperm genes. Candidates for directional selection were defined as the 5% most extreme values (bimodal test) for branch length, Tajima’s D, or the ratio of non-synonymous to synonymous nucleotide diversity (π_NS_/π_S_) in allogamous populations. Under the null hypothesis of no enrichment, imprinted genes should fall into this class with probability 0.05. We therefore tested whether the observed proportion exceeded 5% using independent binomial tests.

Coevolution was assessed using three complementary measures (Pearson’s correlation coefficients for nucleotide diversity π, branch lengths, and Robinson–Foulds (RF) distances, for each pair of PEGs, MEGs, and endosperm genes), and only correlations exceeding thresholds defined from pair of endosperm genes (1st or 99th percentiles) were retained. Finally, we tested if the number of correlations found between PEGs and MEGs and the number of imprinted genes involved were significantly higher than expected from 1,000 random samplings of 29 endosperm genes.

### Imprinted genes are preferentially expressed in the endosperm

Under the kinship theory, both MEGs and PEGs are expected to be disproportionately represented among endosperm-upregulated genes, because imprinting evolves specifically in tissues exposed to parental conflict. We therefore first tested whether the proportion of upregulated MEGs and PEGs exceeded expectations derived from random sampling of the transcriptome. The proportions of MEGs (0.56) and PEGs (0.30) upregulated in the endosperm were both significantly higher than expected from resampling of all genes (p values < 1e-04), and MEGs were more frequently upregulated than PEGs (based on resampling of imprinted genes; p value =0.03).

Because PEGs are generally assumed to promote offspring resource acquisition, while MEGs tend to restrict it, theory predicts higher optimal expression levels for PEGs, followed by MEGs, compared with non-imprinted endosperm genes. We therefore tested whether expression per expressed allele differed among groups after correcting for gene dosage. After adjusting expression for gene copy number (three expressed copies for non-imprinted endosperm genes, two for MEGs, and one for PEGs), both PEGs and MEGs displayed significantly higher expression levels relative to endosperm genes, with 3.18- and 1.71-fold increases, respectively (bootstrap tests; p-values < 1e-03; Fig 1).

**Figure 1:**
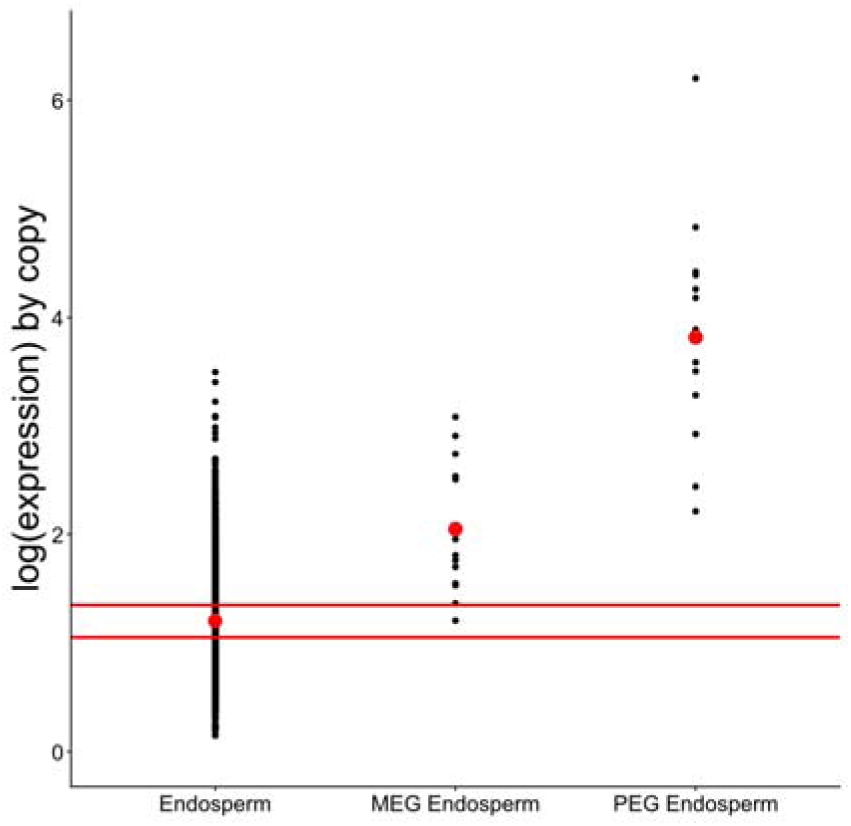
Imprinted genes are associated with higher expression levels in endosperm genes. The dots represent the mean expression level obtains for each gene copy. The red dots represent the mean expression level for each group of genes. The red lines represent 2.5^th^ and 97.5^th^ percentiles of the means obtained for endosperm genes after 1,000 resampling of 29 genes (number of imprinted genes considered in the study). If the observed means of PEG and MEG are outside of these red lines, they are considered significantly different from the mean in endosperm genes, following the bootstrap approach.

Using AIC, we found that the best model to explain this expression pattern (AIC = -4633) was the model that included the gene group, but also the gene size (coefficient = -5.4e-05), the CDS proportion (coefficient = -2.34e-03) and the GC proportion (coefficient = 2.66e-02), whereas the worst model (AIC = -2454) was the one that exclude only the group of genes (AIC = -2454).

We next assessed whether this increased expression was associated with codon or amino acid usage bias. We defined the “overused” and. “underused” codons as codons with a proportion higher and lower respectively than in the 95th percentiles of the 1,000 resample of 29 endosperm genes. Codon and amino acid usage differed significantly between imprinted and endosperm genes, although only two biases were shared by PEGs and MEGs (under-usage of GTC and over-usage of tryptophan; Table S3). However, Predicted High Expression (PHE) scores of codon, derived from overexpressed genes in *A. thaliana* (Sahoo et al., 2019), did not differ significantly across biased and unbiased codons (all p values > 0.09), indicating that codon usage differences were not associated with expression variation.

### The MEGs and PEGs exhibit phylogenetic signals of negative selection in Brassicaceae

To explore phylogenetic patterns, we analysed homologous genes in *A. lyrata* and 21 Brassicaceae species (Table S4). As a proxy for gene turnover, we compared the number of species with homologous sequences across groups. Thus, endosperm genes were conserved across significantly more species than other categories, and PEGs more than control genes (Fig 2; Table S5). The best model to explain this pattern (AIC = 8696) was the model that included the gene group, the gene size (coefficient = 9.74-04) and the GC proportion (coefficient = 0.19), whereas the worst model (AIC = 9104) was the one that exclude the gene size.

**Figure 2:**
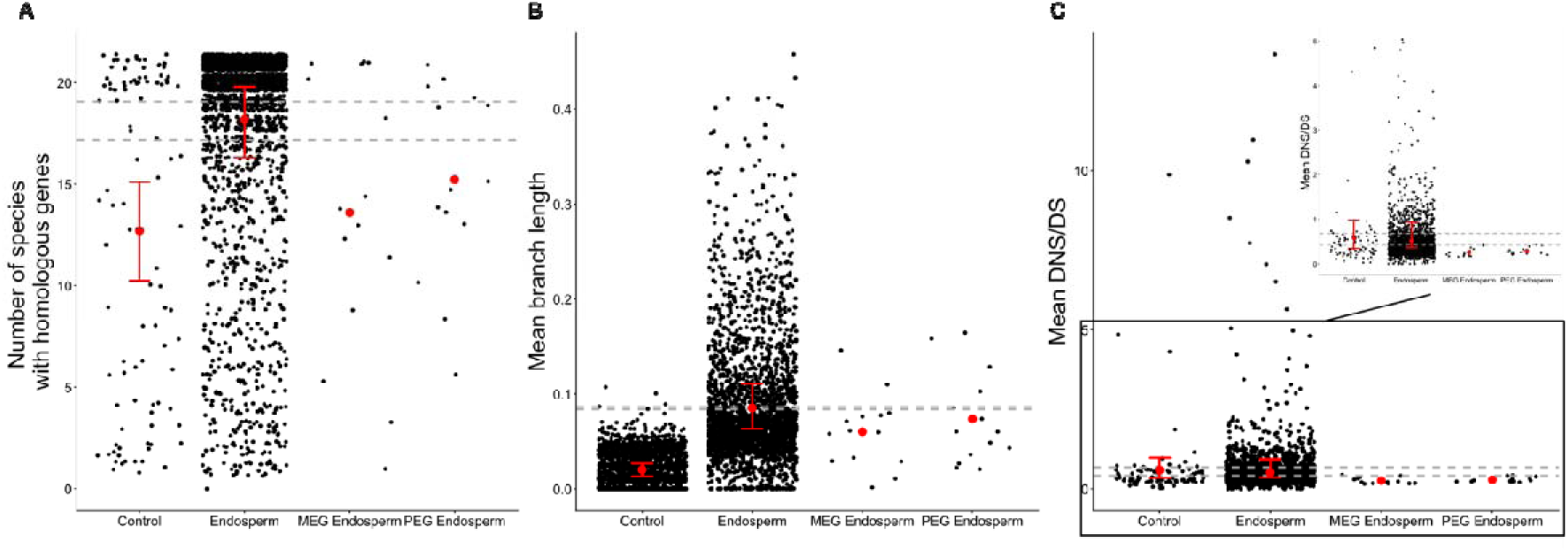
Imprinted genes present phylogenetic signals of negative selection compared to endosperm genes across the Brassicaceae. A: Variations of the number of species with homologous genes by the group of genes. B: Variations of the mean branch length by gene based on the phylogenetic tree obtained by Hasegawa–Kishino–Yano 1985 model based on the 1055 endosperm genes found in all species. C: Variations of the D_NS_/D_S_ by the group of genes with a focus on values lower than 5 (top right). Two extremes values (>200) were removed. The grey dots represent the mean value obtained for one gene. The red dots represent the mean in each group of gene. The dashed lines represent 2.5^th^ and 97.5^th^ percentiles of the means obtained for endosperm genes after 1000 resamples of 100 genes (size of group of control genes). If the real mean of control genes was not between these dashed lines, the value was considered significantly different from that of the endosperm genes, following bootstrap approach. The red lines represent 2.5^th^ and 97.5^th^ percentiles of the means obtained for endosperm and control genes after 1,000 resamples of 29 genes (the number of imprinted genes considered in the study). If the real means of PEG and MEG were not within these red lines, the values were considered significantly different from those in endosperm or control genes, using the bootstrap approach.

Gene-tree topologies of MEGs and PEGs did not differ significantly more from the species tree than endosperm genes (based on RF distances; bootstrap tests; p value = 0.60 for PEGs; p value = 0.34 for MEGs), so branch lengths (proxy for evolutionary rates; Fig. S1) were estimated directly from the species-tree topology for genes having homologous genes in at least three species (including *A. lyrata*). Thus, endosperm and imprinted genes show significantly longer branches than in control genes (Fig 2; bootstrap; Table S5), with significantly longer branch in PEGs and in endosperm genes. The best model to explain this pattern (AIC = -15231) was the model that included the gene group, the gene size (coefficient = 9.73e-06), the CDS proportion (coefficient = -8.10e-04) and the GC proportion (coefficient = 9.63e-04), whereas the worst model (AIC = -14976) was the one that exclude the gene size.

Finally, the mean ratio of non-synonymous to synonymous substitution rate (D_NS_/D_S_) between *A. lyrata* and the other Brassicaceae species decreased significantly in imprinted genes compared to endosperm genes (Table S5), indicating a signal of purifying (negative) selection. The best model to explain this pattern (AIC = 13382) was the model that included the gene group and the GC proportion (coefficient = 0.18), whereas the worst model (AIC = 13401) was the one that exclude the group of genes.

Together, these phylogenetic patterns indicate long-term evolutionary constraints in endosperm genes, a signal that can arise under ancient balancing selection or strong functional conservation. Importantly, these phylogenetic inferences reflect deep evolutionary timescales and therefore cannot be directly compared to the signatures of balancing selection at the population level, which we analyse in the next section.

### Imprinted genes show elevated π_NS_/π_S_ independent of the mating system

To evaluate the effect of the mating system and imprinting on polymorphism, we compared Tajima’s D, π and the π_NS_/π_S_ for imprinted, endosperm and control genes across nine allogamous and nine autogamous populations (Table S6). A previous study reported that the mating system affects seed size through pollen in these populations (Willi, 2013), suggesting a reduced intensity of parental conflict in autogamous populations.

Under directional selection, reductions of Tajima’s D, π and π_NS_/π_S_ are expected, whereas balancing selection is expected to increase these statistics. Because calculations of these statistics require sufficient polymorphism in all populations, genes with fewer than 0.5% polymorphic sites were excluded from Tajima’s D, and genes with π_S_ = 0 from π_NS_/π_S_ analyses, resulting in a significantly higher retention rate for endosperm genes than for control genes for Tajima’s D and π_NS_/π_S_ (binomial tests; Table S7). However, more importantly, these filters retained similar proportions of genes across endosperm and imprinted genes, and across imprinted and control genes (Table S7). Thus, the different signatures of selection detected in Tajima’s D and π_NS_/π_S_ for the imprinted genes, relative to endosperm genes in the allogamous and autogamous populations in the following sections, cannot be attributed to unequal gene retention nor to systematic biases against low-diversity loci, supporting the interpretation that these signals reflect genuine biological differences rather than filtering artefacts.

In allogamous populations, Tajima’s D was significantly higher in endosperm genes than in control genes, while π and π_NS_/π_S_ showed no significant differences. Imprinted genes showed an overall increase in π_NS_/π_S_ relative to non-imprinted endosperm genes (Fig 3; Table S8). The best model to explain this pattern of Tajima’s D (AIC = 9322) was the one that included only the gene size (coefficient = -1.59e-04), whereas the worst model (AIC = 9337) was the one that excluded only the gene size. For the π, the best model (AIC = -36369) was the one that included the gene size (coefficient = -9.89e-08) and the GC percentage (coefficient = -5.59e-05), whereas the worst model (AIC = 9337) was the one that excluded only the gene size. Finally, the best model to explain this pattern of π_NS_/π_S_ (AIC = -721) was the one that included only the proportion of CDS (coefficient = 2.21e-03), whereas the worst model (AIC = 9337) was the one that included all the parameters.

**Figure 3:**
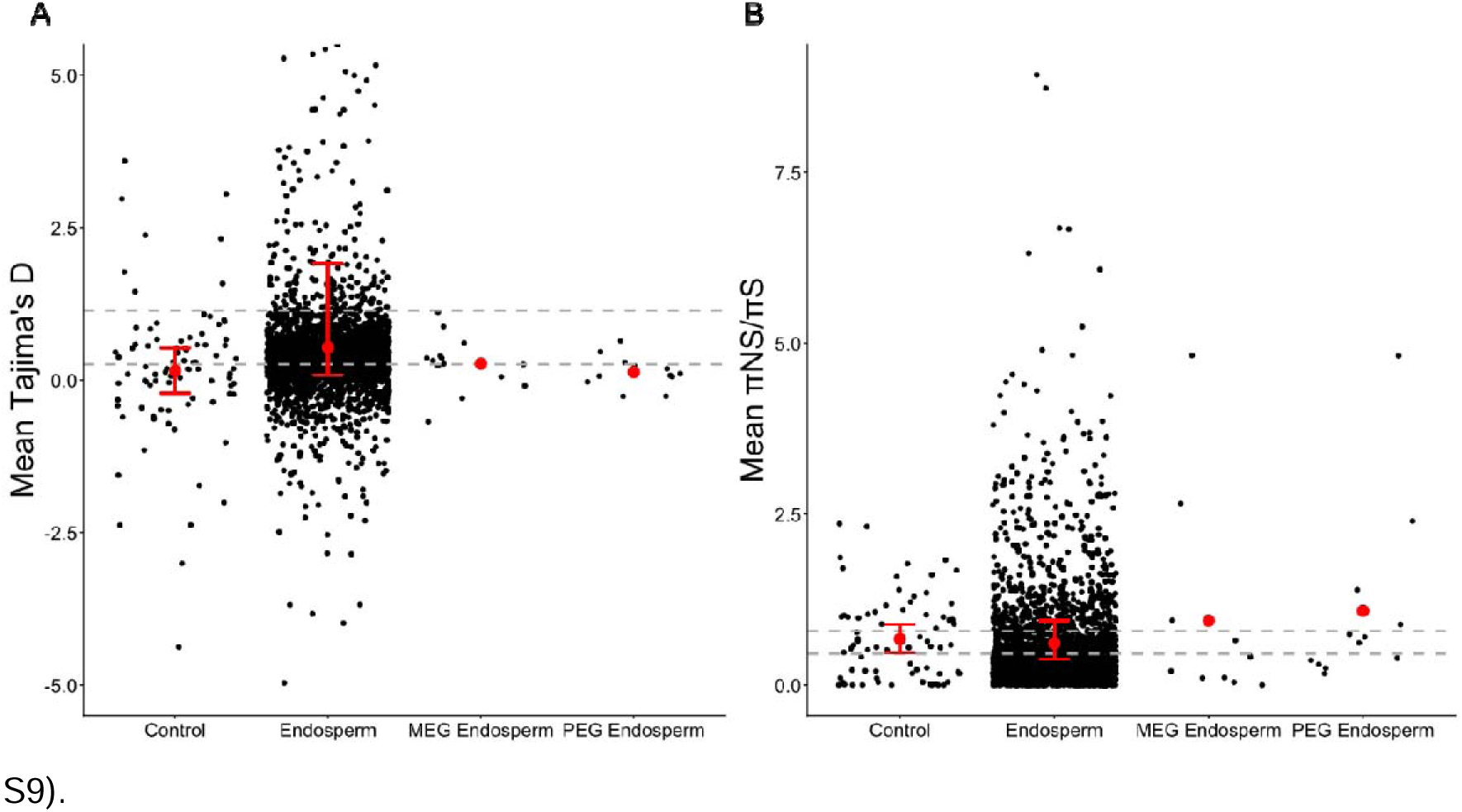
Imprinted genes present an unusual decrease of Tajima’s D associated with an increase of π_NS_/π_S_ in allogamous populations compared to endosperm genes. A: Variations of Tajima’s D among genes group. Only the genes with a mean between -5 and 5 are indicated. B: Variations of the π_NS_/π_S_ among genes group. Only the genes with a mean between 0 and 1.1 are indicated. The grey dots represent the mean value obtained for one gene. The red dots represent the mean of each gene group. The dashed lines represent the 2.5^th^ and 97.5^th^ percentiles of the means obtained for endosperm genes after 1000 resamples of 100 genes (size of the control gene group).. If the real mean of control genes was not between these dashed lines, the value was considered as significantly different than in endosperm genes. The red bars represent the 2.5 and 97.5^th^ percentiles of the means obtained for endosperm and control genes after 1,000 resamples of 29 genes (number of imprinted genes considered in this study). If the real means of PEG and MEG were not between these red lines, the values were considered significantly different from those in endosperm or control genes, following the bootstrap approach.

In autogamous populations, Tajima’s D in endosperm genes no longer differed from controls (Table S8). Moreover, MEGs and endosperm genes exhibited decreased π_NS_/π_S_ compared to control genes (Table S8). These trends in Tajima’s D and π_NS_/π_S_ are explained by the mating system that only significantly affects endosperm genes (Kruskal-Wallis tests; Table S9).

### A limited number of MEGs exhibited strong signals of directional selection

To assess whether loci subject to imprinting show an enrichment in signatures of directional selection, we tested whether they occur more frequently than expected in the extreme tail of the empirical distribution of endosperm genes. Candidates for directional selection were defined as the 5% most extreme values (bimodal test) for branch length, Tajima’s D, or π_NS_/π_S_ in allogamous populations. However, none of these proportions were significantly higher than expected (p = 0.66, 1.00 and 1.00 respectively).

We identified only four MEGs potentially under strong directional selection (Table S1). Two MEGs were associated with a reduced branch length (Table S1): one lacked homologous gene in *A. thaliana*, the second encoded an AGAMOUS-like protein expressed during proliferative endosperm development (Day et al., 2008), involved in female gametophyte, seed and early endosperm development (Bemer et al., 2010; Zhang et al., 2018). One MEG identified by a reduced Tajima’s D encodes a F-box domain–containing protein regulated by non-CG DNA methylation in *A. thaliana* (Garro et al., 2025) and associated with the regulation of stress-responsive genes (Papdi et al., 2008). Finally, one MEG encoding a Dof-type zinc finger protein involved in seed coat formation (Zou et al., 2012) shows a reduced π_NS_/π_S_.

### Clear trend of coevolution among imprinted genes in Brassicaceae

Imprinting could promote coevolution between genes favouring paternal or maternal fitness in an arms race scenario. To test this hypothesis, we investigated potential coevolutionary signals between PEGs, MEGs and endosperm genes, based on (i) nucleotide diversity (π) within populations, (ii) branch length in phylogenetic trees, and (iii) RF distances between tree topologies. Because the coevolution hypothesis is more closely related to noncoding regions, we focused on π rather than π_NS_/π_S_.

First, using thresholds defined from endosperm genes (−0.66 and 3.20 for 1 and 99% of distribution), 13 significant Pearson correlation coefficients involving 6 PEGs and 9 MEGs were detected. Both the number of PEGs-MEGs correlations and the number of imprinted genes involved exceeded expectations (p-values= 4.5e-02 and <1e-03 respectively; Table S10). Considering endosperm genes significantly correlated with these imprinted genes, as well as indirect links between imprinted and endosperm genes, the resulting network contained 14 MEGs, 13 PEGs, and 1,643 endosperm genes (=1,670 genes; Fig 4.A; Table S10). This network did not show enrichment for specific Gene Ontology (GO) terms compared to random networks of equal size. Notably, one PEG encoding a chloroplast-localized protein required for embryonic development (Bryant et al., 2010) was correlated with more genes than 99% of endosperm genes (Table S1 and S10).

**Figure 4:**
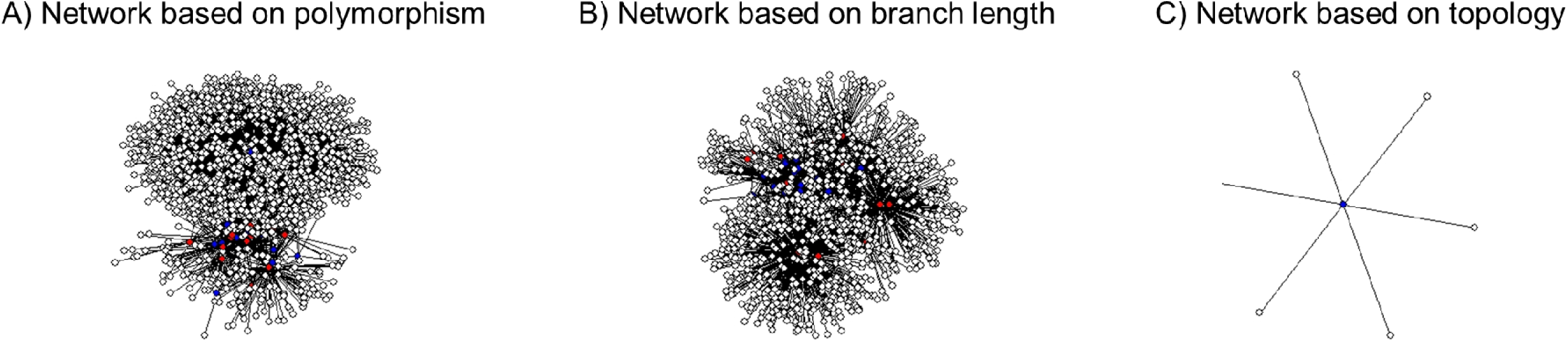
Networks of the imprinted and endosperm genes based on A) polymorphism in population, B) branch length and C) phylogenetic topologies across the Brassicaceae. A) Network based on the Pearson’s correlations of π found on genes in the populations. B) Network based on the Pearson’s correlations of the branch length for each gene estimated with the global topology. C) Networks based on the RF distance of topologies. The black and grey lines represent the negative and positive correlations respectively. Blue dots: PEG. Red dots: MEG. White circles: non-imprinted endosperm genes.

Following the same methodology, the thresholds based on branch length (−0.64 and 0.89) identified 16 significant correlations among 6 PEGs and 9 MEGs. Both the number of PEGs-MEGs correlations and the number of imprinted genes involved exceeded expectations (p-values = 8e-03 and 3e-03 respectively; Table S11). None of these correlations was in common with the previous 13 significant diversity correlations. Considering endosperm genes significantly correlated with these imprinted genes, the resulting network contained 14 MEGs, 14 PEGs, and 1,027 endosperm genes (=1,055 genes; Fig 4.B; Table S11). This network was enriched for the GO categories “*negative regulation of catalytic activity”* and “*syntaxin binding”* (Table S12). One MEG in this network, encoding the AGAMOUS-like protein previously mentioned, was correlated with more genes than 99% of endosperm genes (Table S11).

Finally, we investigated gene pairs that showed high tree similarity (RF = 0) but stronger deviation from the species topology than 95% of endosperm genes (RF > 0.71 with species topology; Fig. S1). Based on this approach, the topology of only one PEG encoding an F-box protein was associated with six endosperm genes (Fig. 4.C; Table S13). This small network was enriched for the GO term *DNA-binding transcription factor activity* (Table S12). Moreover, the PEG exhibited a significantly higher number of associations than 99% of non-imprinted endosperm genes (Table S1; Table S13).

Across all analyses, 22 pairs including 29 imprinted genes were identified. To test for coevolution between specific mutations within these pairs, we applied the clustering approach implemented in the Cosubstitution Mapping (CoMap) package (Dutheil and Galtier, 2007).

Specifically, we searched for groups of 2-10 coevolving sites within each candidate pair of imprinted genes. To limit false positives, the number of such groups detected in candidate pairs was compared with that observed in non-candidate pairs of imprinted genes that were not associated in the previous networks and were present in at least three species. CoMap did not detect an excess of coevolving residues in candidate pairs relative to non-candidate imprinted pairs (Table S14). These directly correlated pairs of imprinted genes were therefore not associated with elevated site-level coevolution.

## Discussion

The kinship theory remains the prominent selective framework for explaining the evolution of genomic imprinting (Patten et al., 2014). According to the kinship theory, imprinting is expected to evolve in contexts of multiple mating where parental genomes have conflicting interests over resource allocation, and alleles at loci subject to genomic imprinting are therefore expected to be disproportionately expressed in the endosperm. The footprint of parental conflict on genome sequences translates into signatures based on sequence (*e.g.,* polymorphism or divergence) is model-dependent and may involve selection acting on regulatory mechanisms rather than coding sequences of loci subject to imprinting. Several models have further proposed that the resolution of parental conflict through imprinting (Wilkins and Haig 2001; Mills and Moore 2004), and related intralocus conflict models, could be associated with directional selection and/or coevolution among antagonistically acting loci (see also Day and Bonduriansky 2004). However, a selection acting primarily on alleles affecting gene expression may attenuate sequence-based signatures, and their sensitivity to mating-system variation remains unclear.

Here, we tested these predictions by comparing imprinted and non-imprinted endosperm genes in *A. lyrata*, an allogamous species that has shifted to autogamy in some populations from North America, and assessing the impact of the mating system shifts. While several figures follow a unified graphical structure to facilitate comparison across gene categories, the underlying analyses encompass expression-level differences, population genetic statistics, phylogenetic metrics, and coevolutionary patterns, providing a broader evaluation of theoretical expectations derived from models of parental conflict and imprinting (Miyake et al., 2009; Hatorangan et al., 2016; Tuteja et al., 2019). Although we detected patterns consistent with predictions of kinship theory and related intralocus conflict models, the signals differed across statistics and were not consistently replicated among analyses. This suggests that selection associated with parental conflict may act heterogeneously across loci or primarily through regulatory rather than coding changes, and underscores the importance of integrating multiple lines of evidence when assessing evolutionary hypotheses of imprinting.

### Patterns compatible with unresolved intralocus conflict in endosperm genes

In the absence of genomic imprinting, genes involved in unresolved intralocus conflict are expected to evolve under balancing selection (Day and Bonduriansky 2004; Sylvestre et al., 2023). In line with this expectation, non-imprinted endosperm genes were more widely conserved across Brassicaceae, suggesting lower turnover, and displayed longer phylogenetic branches (Edwards, 2009; Kasimatis et al., 2017; Llaurens et al., 2017). This interpretation is supported by model comparisons showing that the gene group contributes to explaining variation in these statistics. In addition, these genes showed elevated Tajima’s D values in allogamous populations. However, the absence of a parallel increase in polymorphism, and the absence of this parameter in the best model predicted by AIC, suggests that this signal may not solely reflect balancing selection.

Conflict over resource allocation is predicted only in multiple mating systems (Haig 2000). This parental conflict is expected to be reduced in autogamous systems, which may alter selective regimes acting on endosperm-expressed genes. In line with this, Tajima’s D and π_NS_/π_S_ decreased in these populations, suggesting that reduced parental conflict may be associated with patterns compatible with directional selection and that the effect of the mating system is detectable even in populations that have recently and partially transitioned to autogamy (Ross-Ibarra et al., 2008; Foxe et al., 2010; Hough et al., 2013; Willi et al., 2018).

It is important to emphasize that mainly the phylogenetic analyses, but only Tajima’s D for the analysis at the population level, suggest signals compatible with balancing selection in endosperm genes. Phylogenetic patterns reflect long-term evolutionary processes, potentially including ancient or persistent balancing selection, whereas the statistics at the population level detect more recent forms of balancing selection and are sensitive to demography, recombination, and reduced polymorphism. Therefore, the absence of consistent evidence at the population level does not contradict the phylogenetic signal but rather indicates that any balancing selection acting on these loci is either ancient, weak, or too localized to be detected by gene-wide summary statistics.

Importantly, the absence or inconsistency of the signatures at imprinted loci at the population level does not contradict the existence of parental conflict. Recent models suggest that selection may predominantly target regulatory mechanisms controlling imprinting or expression levels, rather than coding variation at imprinted genes themselves, thereby limiting detectable signatures of selection in analyses based on coding sequence (Flintham et al., 2025).

### Codon usage bias and modulation of expression in imprinted genes

Kinship theory predicts that imprinted genes should be preferentially expressed in the endosperm (Haig 2000; Patten et al., 2014). Consistent with this prediction, imprinted genes were enriched among endosperm-expressed genes. Interestingly, the MEGs are more strongly associated with endosperm expression than PEGs, as observed in maize (Eaux et al., 2013). Yet, more than half were also upregulated in other tissues suggesting that distinct selective pressures unrelated to parental conflict may act on them.

An important debate suggests that detecting directional selection in imprinted genes is difficult because selection may primarily target gene expression (Patten et al., 2014). Consistent with this idea, both PEGs and MEGs showed increased expression per gene copy, and gene group was the factor most frequently retained by AIC-based model selection. Interestingly, the PEGs are associated with increased expression levels, despite gene size reducing expression efficiency globally (Grishkevich and Yanai, 2014), including in our study.

Moreover, imprinted genes displayed codon usage biases relative to endosperm genes, however, we were not able to determine if these biases enhance or inhibit the expression of imprinted genes (Sahoo et al., 2019). More studies with direct analyses of the effect on expression of the different codon usage are required to verify this hypothesis in the future.

### Phylogenetic signals of ancient purifying selection in imprinted genes

Several models related to parental conflict have proposed that the resolution of conflict through imprinting could be associated with directional or purifying selection (Wilkins and Haig 2001; Mills and Moore 2004). In line with this idea, our phylogenetic analyses showed that imprinted genes exhibit lower conservation of homologous genes across Brassicaceae species, likely reflecting a higher turnover driven by accelerated evolutionary rates. Furthermore, imprinted genes displayed reduced D_NS_/D_S_, indicative of strong purifying selection, and the group of genes is the most important parameter in models to predict this pattern.

This phylogenetic signal of purifying selection was also evident in MEGs when branch lengths were compared to those of endosperm genes. We note that two MEGs, including AGAMOUS-like protein implicated in one coevolutionary network, exhibited branch lengths suggesting a particularly strong negative selection. Our observations were not in line with the positive selection detected on PEGs on the autogamous species *A. thaliana* and *C. rubella* (Hatorangan et al., 2016; Tuteja et al., 2019), raising the question of how the mating system impacts the selection of imprinted genes in these species.

Nevertheless, caution is warranted in attributing the observed purifying directional selection solely to imprinting. In fact, the detected purifying selection could be explained by alternative mechanisms (Patten et al., 2014), including (1) an increase of the efficiency of selection by hemizygous expression (Haig and Westoby 1989), (2) selection against Transposable Element (TE; Padilla-García et al., 2025) or (3) local adaptation (Kremer et al., 2025). However, none of the current models rigorously predict the effects of these mechanisms on polymorphism and divergence, making it difficult to discriminate among alternative explanations.

### Implications for reproductive isolation and adaptation

The evolutionary consequences of purifying selection on alleles at imprinted loci remain poorly understood. Directional selection on the coding regions of these genes could (1) influence adaptive potential by shaping the spectrum of selectable alleles within populations (Cvijović et al., 2018), (2) contribute to speciation processes by generating genetic incompatibilities between lineages (Seehausen et al., 2014), (3) facilitate the transition to apomixy in plants (Hojsgaard and Schartl 2021) and (4) promote the establishment of reproductive isolation (Maheshwari and Barbash 2011; Cvijović et al., 2018; İltaş et al., 2021), more particularly through endosperm-based post-zygotic barriers (Haig 2000; Spencer 2000; Kinoshita 2007; Köhler et al., 2010).

Previous studies empirically demonstrated that some imprinted genes are involved in these barriers, including in the autogamous species *A. thaliana* (Wolff et al., 2015), between the allogamous *Solanum peruvianum* et *S. chilense* (Florez-Rueda et al., 2016), between the allogamous *Capsella grandiflora* and the autogamous *C. rubella* and *Capsella orientalis* (Lafon-Placette et al., 2018), between the allogamous *Mimulus guttatus*, *M. luteus* (Kinser et al., 2021), *M. caespitosa* and *M. tilingi* (Sandstedt and Sweigart 2022), and between the autogamous wheat species *Triticum durum* and *T. aestivum* (Yang et al., 2023).

Imprinted genes may also act as central hubs within complex regulatory networks that contribute to hybrid vigour and reproductive barriers (Botet and Keurentjes 2020). However, despite these observations, the broader evolutionary implications of these effects at the network level remain largely unresolved, particularly regarding how they interact with demographic history, variation in the mating system, and other selective pressures.

### Imprinted genes are associated with singular signals of selection in North America

PEGs and MEGs were associated with higher π_NS_/π_S_ values compared to endosperm and control genes, which could indicate balancing or relaxation of selection on coding regions (Galtier and Rousselle 2020). However, this signal was not observed for Tajima’s D. This apparent relaxation may not reflect parental conflict but instead could result from the recent colonization of North America by *A. lyrata* from Europe (Ross-Ibarra et al., 2008; Otto et al., 2015; Willi et al., 2018) that could have reduced effective population size (Galtier and Rousselle 2020) and the efficiency of purifying selection (Laenen et al., 2018), especially on TE (Padilla-García et al., 2025). Testing this hypothesis would require future comparisons of selection signals in endosperm and imprinted genes across allogamous populations with varying levels of polymorphism. Nevertheless, we note that two MEGs exhibited reduced Tajima’s D or π_NS_/π_S_, consistent with ongoing directional selection.

### Imprinted genes are not sensitive to shifts in the mating system in A. lyrata

Conflict over resource allocation is expected to be reduced in autogamous lineages relative to allogamous ones (Haig 2000; Brandvain and Haig 2005). However, most studies of genetic and phylogenetic signatures of selection associated with imprinting have focused on autogamous species, leaving the effect of shifts in the mating system on polymorphism of imprinted genes unresolved. In the autogamous species *A. thaliana*, the imprinted *MEDEA* locus shows increased polymorphism but not in its allogamous sister species *A. lyrata* (Spillane et al., 2007; Miyake et al., 2009). However, PEGs were associated with positive selection in the autogamous species *A. thaliana* and *C. rubella* (Hatorangan et al., 2016; Tuteja et al., 2019). In our study, the selection signals in imprinted genes, contrary to endosperm genes, showed limited sensitivity to variation of the mating system. The absence of variation in Tajima’s D could be explained if directional selection had already fixed the optimal alleles prior to the transition from allogamy to autogamy, which could also explain the signal of positive selection detected in PEGs in *A. thaliana* and *C. rubella* (Hatorangan et al., 2016; Tuteja et al., 2019). However, this explanation does not account for the high π_NS_/π_S_ observed in allogamous and autogamous populations. Theoretical studies are necessary to estimate the combined effect of selection related to the kinship theory, imprinting, demographic history and the mating system.

### Imprinted genes present signals of ancient and recent coevolution

Another potential, but not mandatory, outcome proposed in extensions of the kinship theory is the coevolution of some imprinted loci with antagonistic effects on offspring growth (Moore and Haig, 1991; Wilkins and Haig 2001; Mills and Moore, 2004). To investigate this, we applied both population genomic and phylogenetic approaches (Dutheil and Galtier 2007). We found that most PEGs and MEGs were connected within coevolutionary networks, with a significantly higher number of direct associations than expected by chance. These networks encompass essential functions for endosperm development.

For instance, one network based on branch length correlations was enriched in enzymatic activities related to cell wall formation and modification, a key process during endosperm cellularization in Arabidopsis (Hehenberger et al., 2012), which in turn influences seed size (Pignatta et al., 2018; Zhang et al., 2020), a likely target of parental conflict. These correlated evolutionary rates could reflect equivalent selective constraints (Galtier and Rousselle 2020) or just a common chromosomal context (*e.g.* TE) that modifies the mutation rate (Lynch 2010). Interestingly, this network appeared to be strongly dependent on a MEG, an AGAMOUS-like protein, characterised also by reduced branch lengths, consistent with expectations that coevolution may drive coordinated evolution among genes primarily through rare compensatory mutations (Fraser et al., 2002)

The network based on polymorphism involved genes related to syntaxin-binding proteins, crucial for membrane activity, molecular transport, and protein storage, whereas the topology-based network was enriched in DNA-binding transcription factors, including those regulating seed development, tissue differentiation, and storage metabolism (Noguero et al., 2013).

These phylogenetic and population genetic signals of coevolution could promote the maintenance of the gene functions (Clark et al., 2012), limit their evolutionary independency (Yang and Bielawski 2000) and be another source of reproductive barrier (Presgraves 2010). Interestingly, the coevolution signal did not involve increased site-by-site coevolution, suggesting that protein–protein molecular interactions, often drivers of coevolution (Green et al., 2021), played a limited role here. Instead, the observed patterns may reflect selection acting on gene dosage rather than protein products (Clark et al., 2012).

However, as correlations between evolutionary statistics can arise from variation of the mutation rate, recombination environment or shared functional constraints, we interpret them as exploratory signals only. The more stringent CoMap analysis (Dutheil & Galtier 2007) did not reveal enrichment for co-evolution at the site level, indicating that networks based on correlation should be treated cautiously and do not by themselves confirm co-evolution.

### Main limitations in our study

Although our study combines multiple phylogenetic and population genomic statistics, the signals detected were not consistently congruent across approaches. This inconsistency may reflect evolutionary processes not accounted for, (such as demographic history or chromosomal context) as well as several methodological limitations, outlined below.

First of all, estimating Tajima’s D from pooled sequencing data is challenging because statistics based on the Site Frequency Spectrum (SFS) are sensitive to coverage variation and sampling noise, which increase estimation variance (Futschik & Schlötterer 2010; Ferretti et al. 2013; Lynch et al. 2014). Although we minimized biases specific to the dataset by comparing genes processed under identical conditions, technical noise cannot be fully excluded. In addition, short genes, such as many MEGs or control genes, yield fewer segregating sites, reducing the reliability of Tajima’s D (Tajima 1989).

A second limitation concerns the imprinting dataset. The set of imprinted genes analysed was derived from a study on a geographically and genetically distinct allogamous subspecies European *A. lyrata petraea* (Klosinska et al., 2016), assuming conservation of imprinting, because no within-lineage data on imprinting in *A. lyrata* exist. However, imprinting shows limited conservation within *A. thaliana* (Gehring et al., 2011; Wolff et al., 2011; Hsieh et al., 2011; Pignatta et al., 2014), between *A. thaliana* and *A. lyrata* in our data, and generally across species (Eaux et al., 2013; Hatorangan et al., 2016; Roth et al., 2018; Flores-Vergara et al., 2020). It is therefore possible that several genes classified as imprinted in our analyses are not imprinted in the studied populations, especially given known population-level variation in imprinting drive by TE (Padilla-García et al., 2025). Future population-specific characterisations of imprinting will be critical to resolve this issue.

Moreover, some imprinted genes identified by Klosinska et al. (2016) may have been inadvertently excluded from our dataset due to our method for detecting tissue-specific expression, which relied on RNA sequencing (RNA-seq) data pooled from both allogamous and autogamous populations. More controlled and population-specific experimental designs will be required to recover the complete set of endosperm-expressed imprinted genes relevant to parental conflict.

A last important limitation is about the panels of genes compared (control, endosperm and imprinting genes). To be consistent with previous studies (Spillane et al., 2007; Miyake et al., 2009; Hatorangan et al., 2016; Tuteja et al., 2019), we also selected genes from panels based on expression patterns. However, the three gene panels differ substantially in gene size, CDS proportion, and GC content, factors known to influence both polymorphism and phylogenetic metrics, but seldom considered in previous studies (Spillane et al., 2007; Miyake et al., 2009; Hatorangan et al., 2016; Tuteja et al., 2019).

Endosperm genes and PEGs are substantially longer, whereas MEGs and control genes are shorter. The size of the gene is an important parameter that impact many statistics. For example, larger genes tend to evolve more slowly due to increased functional constraint and pleiotropy (Massey 2013; Lopes et al., 2021), display higher conservation and reduced turnover (Abdel-Haleem 2007; Koonin 2015), and can inflate the number of species with detectable homologous genes. In this sense, we found a positive correlation between the number of species with homologous genes and the gene size. This higher gene size may partially explain the higher homology counts observed for PEGs and endosperm genes relative to control genes. Conversely, we found that the gene size is negatively correlated with polymorphism and Tajima’s D. However, this tendency is in contradiction with longer genes as endosperm genes presenting a higher Tajima’s D or shorter genes as the MEGs being associated with lower polymorphism, suggesting that, maybe, the effects of parental conflict on polymorphism in these genes are underestimated, more specifically for MEG because shorter genes may yield less stable estimates of Tajima’s D (Tajima 1989). We found also a negative correlation with the expression level, that could explain partially why the expression of MEG is more efficient, even if the gene group stays the most redundant parameter predicted by AIC, but could not explain why the expression of PEG is higher than endosperm genes.

Several studies show that long, constrained genes tend to exhibit shorter branches due to stronger purifying selection (Kimura 1983; Abdel-Haleem 2007; Massey 2013; Koonin 2015). However, in our study, we found a positive correlation between the length of the branches and the gene size, maybe explained by the amount of endosperm genes with long branches. This correlation suggests that the longer branches observed in imprinted and endosperm genes are not explained by the size of the genes.

Moreover, both imprinted and endosperm genes show elevated CDS content relative to controls. Thus, this parameter could not explain the increase in efficiency observed in expression of imprinted genes, despite the negative correlation detected. In the same way, this parameter could not explain the reduced π_NS_/π_S_ observed in imprinted genes compared to endosperm genes, despite the positive correlation detected. Interestingly, high CDS proportion is associated with stronger purifying selection (Hughes et al., 2003), which can reduced the length of the branches, as observed in our study. However, the increase of branch length detected in endosperm and imprinted genes is the reverse pattern predicted, suggesting that the effect on branch length in these genes is maybe underestimated.

Finally, loci subject to imprinting show reduced GC content relative to endosperm and control genes. GC-poor genes tend to have reduced recombination rates, lower polymorphism (Wright et al., 2003), and higher AT-biased substitution rates, which can artificially lengthen branches (Duret & Galtier 2009), and explain the longer branches observed in PEG and endosperm genes. However, contrary to these expectations, we found a negative correlation between the length of the branches and the GC content, which may be explained by the important amount of endosperm genes. Interestingly, we also found a negative correlation between the polymorphism and the GC content in our study, which suggests that the reduced polymorphism observed in MEGs is maybe underestimated. Low GC also reduces the likelihood of finding conserved homologous genes across species (Urrutia & Hurst 2003), as found in our study, but is not able to explain why the PEGs and the endosperm genes are associated with higher number of species with homologous genes despite lower GC content than in control genes. Low GC is expected to also inflate D_NS_/D_S_ by increasing the probability of non-synonymous changes (Kryazhimskiy & Plotkin 2008). However, we detected the reverse correlation in our study, which could explain why imprinted genes are associated with a reduced D_NS_/D_S_.

The lower GC of imprinted genes is also unexpected given their elevated expression (Sharp & Li 1987; Bernardi 2000), suggesting that expression differences may be underestimated. For these reasons, although the three panels differ in genomic features, endosperm-upregulated genes remain the only biologically appropriate baseline for testing hypotheses related to imprinting. A feature-matched control panel would reduce variation but would not provide a more relevant comparison for detecting signatures associated with parental conflict.

Together, differences in gene size, CDS proportion, and GC content may partly account for the heterogeneity of evolutionary signals observed across statistics. Some signatures consistent with parental conflict may therefore be underestimated or confounded, while others may reflect intrinsic genomic properties rather than selective pressures related to imprinting. Future work will require gene panels matched not only for expression, but also for fine-scale genomic features.

Although several genomic features differ among gene panels, we performed targeted robustness checks to ensure that competing non-selective scenarios did not spuriously generate the patterns we report. In particular, the SNP-density threshold retained comparable fractions of MEGs, PEGs, endosperm genes and controls, ruling out retention bias as a driver of polymorphism contrasts. Moreover, all evolutionary inferences were tested against empirical null distributions from endosperm-upregulated genes, which share expression patterns and chromatin context with imprinted genes, limiting confounding effects of gene architecture or local mutation rate. These checks do not rule out all alternative explanations, but they demonstrate that the signals observed are unlikely to arise solely from genomic artefacts or filtering procedures.

### Conclusions

Overall, our analyses revealed that endosperm-expressed and imprinted genes in *A. lyrata* exhibit evolutionary signatures partly consistent with theoretical expectations derived from models of parental conflict and imprinting, including evidence for ancient balancing selection in endosperm genes, expression-level regulation in imprinted genes, ancient purifying selection, and coevolutionary interactions. However, signals at the population level were less consistent with theoretical expectations, likely reflecting demographic history, mating system shifts, and the poor conservation of imprinting across populations. Future work should combine empirical surveys of imprinting with theoretical models explicitly incorporating demography and mating system variation to refine predictions about the evolutionary dynamics of loci subject to genomic imprinting.

## Materials and Methods

We hypothesized, based on the kinship theory, that imprinted genes should display distinct genomic signatures of selection compared with non-imprinted endosperm genes (Table 1). To test this prediction, we designed an approach combining phylogenetics and population genetics. Specifically, we focused on a set of 27 maternally expressed genes (MEGs) and 46 paternally expressed genes (PEGs) previously characterised in *A. lyrata* (Klosinska et al., 2016) and unambiguously annotated in the reference genome V1.0.23 (Hu et al., 2011; https://doi.org/10.5281/zenodo.17287897). These imprinted genes were systematically compared against two control datasets: (i) 2,703 non-imprinted genes upregulated in the endosperm, and (ii) a reference panel of 100 randomly selected ubiquitous genes, expressed across multiple tissues. This design allowed us to disentangle signatures of selection specifically associated with imprinting from those affecting the whole genome or endosperm genes in general.

**Table 1:**
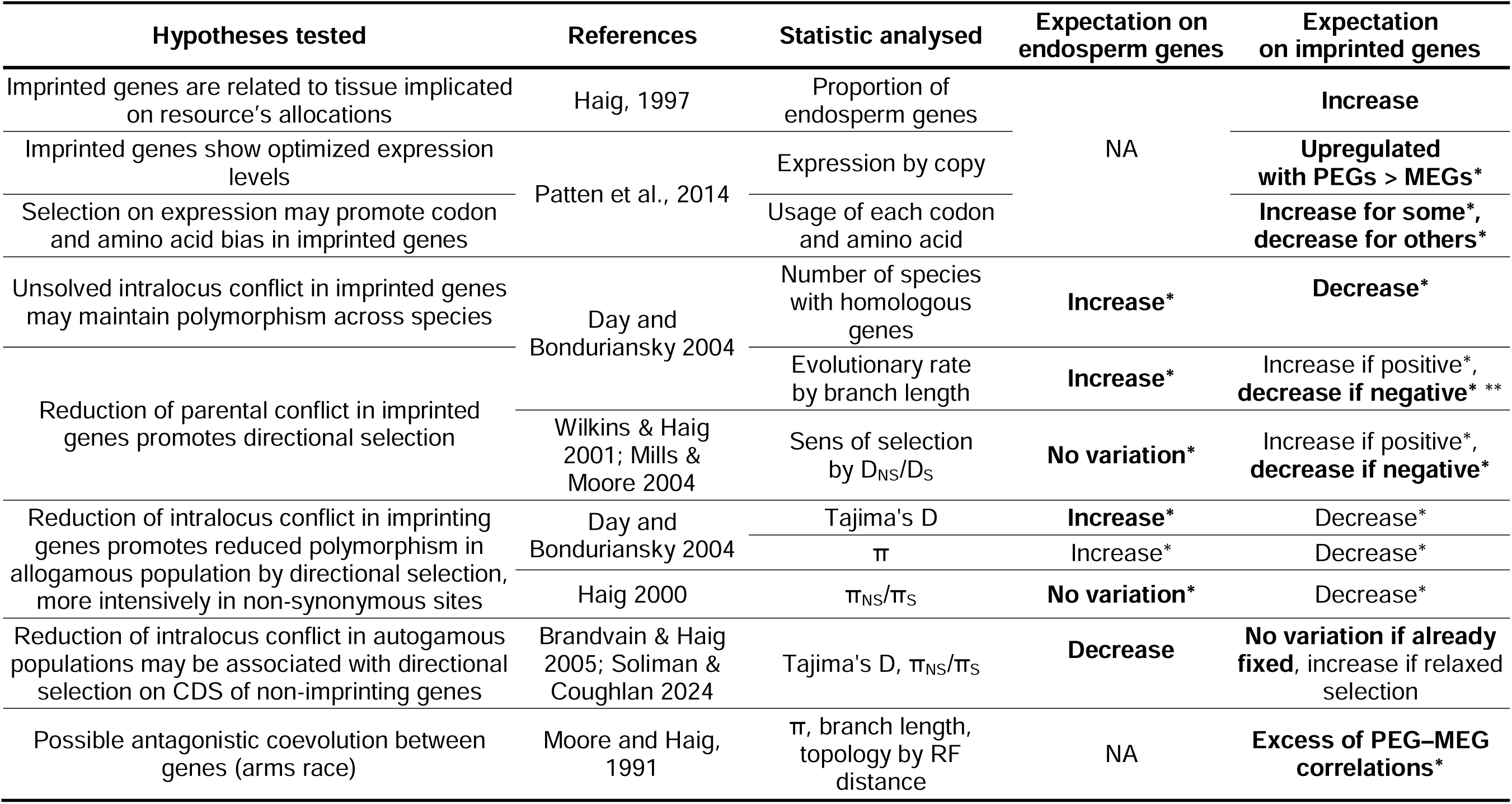
Theoretical expectations derived from models of parental conflict and imprinting on endosperm and imprinted genes and results obtained. Expectations regarding polymorphism, divergence, or coevolution are not explicit predictions of the kinship theory itself, but derive from related models of intralocus or parental conflict and depend on whether selection primarily targets regulatory or coding regions. The expectations validated by the results are signified in bold.* : expectations on endosperm and imprinted genes tested by bootstrap relatively to control and endosperm genes respectively. ** : validated only in MEGs.

### Expression data

RNA-seq expression data from multiple tissues, including leaf, root, pistil, and pollen, were obtained from two *A. lyrata* populations representing 27,820 genes from both European and North American lineages, as described in İltaş et al. (2024). To maximize gene representation and increase the statistical power, the data from both mating type was pooled. To complement these datasets, we also incorporated endosperm RNA-Seq data derived from five allogamous *A. lyrata* individuals, previously generated by Klosinska et al. (2016). The endosperm data were reprocessed and integrated into our analytical pipeline following the instructions and R script available in GitHub (https://github.com/thelovelab/DESeq2). To identify endosperm-specific genes, we retained only those that showed significant upregulation (log2FC > 1, adjusted P < 0.05) in all pairwise comparisons against other tissues. The RNA-Seq datasets for leaf, root, pistil, and pollen are available in the NCBI BioProject database under accession PRJNA1067428, while the endosperm RNA-Seq dataset is accessible under accession GSE76076.

### Genes clusters

We used the list of the 49 PEGs and 35 MEGs found in *A. lyrata* (Klosinska et al., 2016). We kept the genes having conserved coordinates in the reference genome of *A. lyrata* (Hu et al., 2011), resulting in a final list of 27 MEGs and 46 PEGs.

To disentangle signatures of selection specifically associated with the endosperm (where conflicts over resource allocation are expected) from those potentially arising through other selective processes, we separated imprinted genes upregulated in the endosperm (14 PEGs and 15 MEGs) from imprinted genes not showing endosperm-specific upregulation (32 PEGs and 12 MEGs), as determined by differential expression analysis.

To evaluate whether the observed proportions of MEGs and PEGs upregulated in the endosperm deviated from theoretical expectations under the kinship theory, we compared them with null distributions generated by 10,000 random sampling of 49 and 35 genes (the size of the PEGs and MEGs groups) among the 27,820 expressed genes.

For all subsequent analyses, we focused exclusively on endosperm-upregulated imprinted genes and compared them with two reference groups: the set of 2,703 non-imprinted endosperm-upregulated genes (hereafter, “endosperm genes”), the sets of 15 MEG and 14 PEG upregulated in endosperm (hereafter, “MEG” and “PEG”) and a set of 100 randomly selected ubiquitously expressed genes (hereafter, “control genes”). To remain consistent with previous studies on imprinting and parental conflict (Spillane et al., 2007; Miyake et al., 2009; Hatorangan et al., 2016; Tuteja et al., 2019), the control panel was defined based solely on expression patterns. This set is intended to represent a genome-wide baseline that is unrelated to endosperm-specific functions or parental conflict, rather than a genomic feature–matched control. Consequently, no constraints were imposed on gene size, GC content, CDS proportion, or chromosomal position. However, for each statistic studied below, we compared the full model, which included gene group, gene length, CDS proportion, and GC content as explanatory variables, using Akaike’s Information Criterion (AIC). We then evaluated a series of reduced models, removing one parameter at a time, and selected the model with the lowest AIC. This procedure was iterated until the best-supported model was identified using the function StepAIC of the MASS library (version 7.3-65) in R. This approach allowed us to determine (1) whether the gene group remained a significant predictor in the best model and (2) whether genomic features accounted for the observed trends in the analysed statistics.

To test whether expression levels conformed to predictions of higher optimal expression in PEGs than in MEGs, we normalized expression per gene copy number and compared observed values with bootstrap distributions from non-imprinted endosperm genes (10,000 resamples of 29 genes). To account for gene dosage, expression values were normalized by the expected number of expressed copies: three for non-imprinted endosperm genes, two for MEGs, and one for PEGs.

Based on the sequences from the reference genome of *A. lyrata* (Hu et al., 2011), we conducted bootstrap analyses to compare the gene size, the GC and the CDS content (%) of PEGs and MEGs with values obtained from 10,000 resamples of 29 endosperm genes and control genes independently. Similarly, we also conducted bootstrap analyses to compare the values of control genes with those obtained from 10,000 resamples of 100 endosperm genes.

### Estimation of codon and amino acid bias

Based on the sequences from the reference genome of *A. lyrata* (Hu et al., 2011), we estimated the proportion of each amino acid within the gene and the proportion of each possible codon for each amino acid. Then, we conducted bootstrap analyses to compare these proportions in PEGs and MEGs with values obtained from 10,000 resamples of 29 endosperm genes. Thus, we defined the “overused” (resp. “underused”) codons as codons with a proportion higher (resp. lower) than in the 95^th^ percentiles of the 1,000 resample of 29 endosperm genes.

To test if the overused or underused codons were related to gene expression, we compared the variations of Predicted High Expression (PHE) score induced by codons with the score of unbiased codons by permutations test with 1,000 replications for MEGs and PEGs independently. The PHE score was estimated based on overexpressed genes in *A. thaliana* and reflect the importance of a codon on genes expression (Sahoo et al,. 2019). We hypothesized that codons overused in imprinted genes were associated with increases in PHE score because they were associated with increases in gene expression, whereas the underused codons were associated with reduced PHE scores.

### Distribution of the genes across Brassicaceae

We assessed whether the imprinted genes presented specific signals of selection compared to the endosperm and control genes in the phylogeny of 22 Brassicaceae. All the complete sequence of reference genomes were extracted from the Phytozome database (Goodstein et al., 2012; Table S4). We extracted the sequences of the genes in the *A. lyrata* reference genome (Hu et al., 2011) using bedtools V2.30 (Quinlan et al., 2010). Then, we retained the best hit for these sequences against 22 other Brassicaceae using BLAST V2.10 (Altschul et al., 1990) with an e-value < 1e-5. If more than one hit was found for a gene in one species, we filtered the best hit as the hit with the highest length alignment, the highest % of identity, then the highest coverage on the gene in *A. lyrata* and the highest reciprocal coverage on the gene in the other species. Then we extracted the sequence of the best hit for each species of each gene.

To test for significant differences in the number of species with homologous genes across the genes group, we conducted bootstrap analyses to compare the value on PEGs and MEGs with values obtained from 10,000 resamples of 29 endosperm genes and control genes independently. Similarly, we conducted bootstrap analyses to compare the control genes values with those obtained from 10,000 resamples of 100 endosperm genes.

### Divergence of the imprinted genes across Brassicaceae

The rates of nonsynonymous mutations (D_NS_) and synonymous mutations (D_S_) are assumed to follow the nearly-neutral evolutionary process and the ratio D_NS_/D_S_ is therefore approximate to the selective pressure on the protein product of a gene. A decrease in D_NS_/D_S_ in genes indicates a increase of intensity of negative selection. To test whether imprinted genes display accelerated evolutionary rates compared to endosperm genes, using the annotation of the 0-fold and 4-fold degenerate sites in *A. lyrata*, we estimated for each gene the mean evolutionary rates D_NS_/D_S_ between the reference genome of *A. lyrata* and all the species with an homologous gene found. Only the genes found in at least 3 species and with a D_S_ higher than 0 were considered.

To test for significant differences in the D_NS_/D_S_ across the group of genes, we conducted bootstrap analyses to compare the value on PEGs and MEGs with values obtained from 10,000 resamples of 29 endosperm genes and control genes independently. Similarly, we conducted bootstrap analyses to compare the values on control genes with values obtained from 10,000 resamples of 100 endosperm genes.

### Topology of the genes across Brassicaceae

We aligned each gene with all its best hits found across the different species with Muscle V3.8.155 (Edgar 2004). Then, we built a phylogenetic tree for each gene using the HKY85 model with PhyML V3.3 (Guindon et al., 2010). In parallel, we concatenated the alignment of the 1,055 endosperm genes found in the 22 others Brassicaceae to extract the topology of the phylogenetic tree of the endosperm genes build using PhyML V3.3 (Guindon et al., 2010). This tree was used as a proxy of the topology of the endosperm. Then, we estimated the Robinson-Foulds (RF) distance between the topology of this tree and the topology of the tree of each gene obtained previously using the R package TreeDist. Then, to test for significant differences in RF distances across the group of genes, we conducted bootstrap analyses to compare the values obtained from PEGs and MEGs with those from 10,000 resamples of 29 endosperm genes and control genes independently. Similarly, we conducted bootstrap analyses to compare the values on control genes with values obtained from 10,000 resamples of 100 endosperm genes.

### Variation of the branch length across Brassicaceae

We investigated whether the imprinted genes presented variation in the branch lengths compared to the endosperm and control genes. For each gene, we re-estimate gene specific branch lengths using the topology of the phylogenetic tree obtained with the 1,055 endosperm genes using PhyML V3.3 (Guindon et al., 2010). Then, we extracted the terminal branch lengths at the tips for each gene with the R package ‘ape’ version 5.7-1. If the gene was not present in a species, the value was replaced by “NA”. To test for significant differences in the branch length across the group of genes, we conducted bootstrap analyses to compare the mean in PEGs and MEGs with means obtained from 10,000 resamples of 29 endosperm genes and control genes independently. Similarly, we conducted bootstrap analyses to compare the means of control genes with values obtained from 10,000 resamples of 100 endosperm genes.

### Read mapping and variant calling

We used natural accessions of *A. lyrata* from North America from nine allogamous and nine autogamous populations of 25 individuals each (Table S6), using available pool-sequencing data (Willi et al., 2018).

The raw reads from Willi et al (2018) datasets were mapped onto the complete *A. lyrata* reference genome V1.0.23 (Hu et al., 2011) using Bowtie2 v2.4.1 (Langmead and Salzberg 2012). The resulting alignments were then converted to BAM files using samtools v1.3.1 (Li et al., 2009) and duplicated reads were removed with the MarkDuplicates program of picard-tools v1.119 (http://broadinstitute.github.io/picard). Biallelic SNPs in these regions were called using the Genome Analysis Toolkit v. 3.8 (GATK, DePristo et al., 2011) and stored in a GVCF file to keep non-variable called positions. A quality score threshold of 60 was applied using vcftools v0.1.15 (Danecek et al., 2011), to retain only highly reliable SNPs. This stringent cutoff is commonly used in high-coverage datasets to minimise false positives, which is particularly important for downstream analyses of selection and co-evolution that are sensitive to SNP-calling errors (Schlötterer et al., 2014; GATK Best Practices). We excluded sites with less than 15 reads aligned. Sites covered by at least 15 reads but containing no SNP were considered monomorphic. The final number of sites in each sample set is summarised in Table S6.

### Effect of imprinting on global population polymorphism

We decomposed the signal of selection across the imprinted genes into two elementary statistics: Tajima’s D and the ratio of average pairwise heterozygosity between the non-synonymous and synonymous sites π_NS_/π_S_. For the allogamous and autogamous populations, we compared the values obtained for the imprinted genes with those obtained for the endosperm and control.

For each gene of each population, we estimated π, π_NS_/π_S_ for all positions and the Tajima’s D based on allelic frequencies in naturals populations. Briefly, we estimate the Tajima’s D by gene following the classical equation described in Tajima (1989): 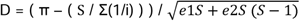, with e1 = (n+1) /3 * (n+1) - (1/ Σ(1/i))/ Σ(1/i) and e2= (2 * (n * n + n + 3)) / (9 * n * (n-1)) - ((n+2) /(Σ(1/i) *n-1)) + (Σ(1/i²) / (Σ(1/i)²)) / (Σ(1/i)² + Σ(1/i²)). S is for the number of segregating sites, i for value between 1 and n-1 haplotypes and n is for the number of haplotypes (number of individuals multiplied by two for diploid).

If a position in the *A. lyrata* genome was covered but not polymorphic, π was set to 0. Then, the 0-fold and 4-fold degenerated sites were identified and extracted from the reference genome and the genome annotation using the script NewAnnotateRef.py (Williamson et al., 2014; updated version available in https://github.com/leveveaudrey/phylogenetic_analyse_brassicacee). Because all mutations at 0-fold degenerated sites alter the sequence of the encoded protein, they are non-synonymous (NS). In contrast, mutations at the 4-fold degenerate sites never alter the encoded amino acid, and all are synonymous (S). For the sake of simplicity, we discarded mutations at 2- or 3-fold degenerate sites, which can be either synonymous or non-synonymous. Based on π estimated for each site, we estimated the mean π_NS_ and π_S_, for each gene in each population, in order to estimate π_NS_/π_S_. We removed the genes with a π_S_ equal to 0. For the Tajima’s D, we applied a SNP-density threshold (≥0.5% polymorphic sites) to ensure reliable estimates. This density-based criterion avoids the gene-length biases inherent in minimum-SNP thresholds, since genes vary widely in size across categories. We verified that this filtering did not disproportionately exclude any gene category: the proportions of retained genes were comparable across endosperm and imprinting genes and across imprinting and control genes (Table S7).

To test for significant differences across the genes group, we conducted bootstrap analyses to compare the mean π, π_NS_/π_S_ and Tajima’s D of PEGs and MEGs with means obtained from 10,000 resamples of 29 endosperm genes and control genes independently. Similarly, we conducted bootstrap analyses to compare the control genes values with those obtained from 10,000 resamples of 100 endosperm genes. We performed the test on allogamous and autogamous populations independently. Finally, for each statistic and group independently, we tested for the mating system effect by a performing Kruskal-Wallis test as implemented in R version 4.3.2.

### Genes in coevolution

To detect coevolving coding sequences, we used three different methods based on the following assumptions: coevolution led BY 1) related polymorphic patterns between genes in populations, 2) related evolutionary rate across species, or 3) similarity in the topologies across species (Dutheil and Galtier 2007).

Correlations between evolutionary rates or diversity metrics were used as exploratory indicators of evolutionary covariance, following approaches such as evolutionary rate covariation, which has been widely applied to detect co-functionality or shared selective pressures (Clark et al., 2012). Because such correlations may reflect genomic or mutational confounders, we used endosperm-upregulated genes as an empirical baseline for defining extreme thresholds.

To detect coevolution based on polymorphism in population and on evolutionary rates across species, we estimated Pearson’s linear correlations between each pair of imprinted and endosperm genes for the mean π and the branch length on each species. Based on these correlations matrices, we extracted correlations between pairs of endosperm genes to define thresholds for extreme values in the distributions. Then, all pairs of genes that included imprinted genes with correlations below the 1% or above the 99% quantiles of the distribution were considered significant. We used these significant correlations of π and branch length to build the coevolution networks.

To detect coevolution based on tree-topology similarity, we first used the distribution of RF distances between endosperm genes and the global endosperm topology to estimate the value obtained in 95^th^ percentile value. Then, we excluded all the imprinted and endosperm genes having a lower RF distance than this value to remove genes whose topology similarity was explained by congruence with the global topology. Finally, we compared the topology of each remaining imprinted gene and endosperm gene. All pairs of genes with a null RF distance were considered as significantly similar and used to build the last coevolution network.

For each network of coevolution, we compared the number of pairs of imprinted genes directly correlated with the number found between genes in 1,000 random samples of 29 endosperm genes. Moreover, we tested for gene enrichment on each GO term in each network based on the occurrences of each term across 1,000 random samples of equivalent size of all the endosperm and imprinted genes. The complete list of GO term associated with each gene was downloaded from Gene Ontology online platform for *A. thaliana* (Ashburner et al,. 2000).

### Identification of coevolving sites

We used a model-based approach to detect coevolving sites for gene pair that showed a coevolution signal. The pairs of imprinted genes studied were all directly associated in at least one previously mentioned network. To use this method, we concatenated each sequence of pairs of genes for each species. If one species was not found in the other gene, the sequence associated with this species was removed.

Coevolving sites within each pair of genes were predicted using the clustering method implemented in the CoMap package (version 1.5.2; https://jydu.github.io/comap/; Dutheil and Galtier 2007). The built-in algorithm was then used to correct for multiple testing and to estimate the false discovery rate (FDR). In this study, 1,000 simulations per tested pair were performed, and the groups of sites with an adjusted p value < 1% were considered coevolving. Simple correlation was used as a measure of coevolution. For each comparison pair, we extracted the coevolving groups and classified them by size (between 2 and 10).

To test for significant excess of correlations, we compared the results obtained for the pair of imprinted genes found in networks previously mentioned with the results obtained for equivalent number of pair comparisons of imprinted genes not associated in any network, for each cluster of coevolving groups of substitutions of equivalent size by Kruskal-Wallis tests. If the number of correlations is significantly higher than the number expected between two random genes by chance or by alignment errors, we expect a significant increase in these numbers in pairs of associated imprinted genes.

## Data availability

The RNAseq data for leaf, root, pistil, and pollen tissues can be found in the NCBI BioProject database under submission number PRJNA1067428. The endosperm RNAseq data is available under submission number GSE76076.

The version of *A. lyrata* reference genome (Hu et al., 2011) used in this paper is available in zenodo (https://doi.org/10.5281/zenodo.17287897).

We used several GitHub repositories, each corresponding to a distinct analytical module. This structure reflects the fact that the datasets and workflows differ substantially across analyses and that several scripts were developed independently and used in other publications. For clarity, all repositories are indexed in a Zenodo record (https://doi.org/10.5281/zenodo.17954040), and each includes documentation, test data, and commented scripts. This multi-repository organization was reviewed and approved for reproducibility and FAIR compliance by the PCI Ecology Data Editor (Checklist PCI Ecology 867).

The R code for the bootstrap tests is available in GitHub (https://github.com/leveveaudrey/bootstrap; https://doi.org/10.5281/zenodo.19051945).

The R code used for permutation test was available in GitHub (https://github.com/leveveaudrey/permutation-test-; https://doi.org/10.5281/zenodo.19051961)

The python code to estimate the proportion of each amino acid within the gene and the proportion of each possible codon for each amino acid is available in GitHub (https://github.com/leveveaudrey/codon-bias; https://doi.org/10.5281/zenodo.19051978).

The python code for the pipeline of phylogenetic analysis (distribution, divergence extraction of topologies and branch length of the genes across Brassicaceae) is available in GitHub (https://github.com/leveveaudrey/phylogenetic_analyse_brassicacee; https://doi.org/10.5281/zenodo.19051955).

The python code for the pipeline including alignment of reads and the SNP calling is available in Github (https://github.com/leveveaudrey/alignment-and-snp-calling; https://doi.org/10.5281/zenodo.19051865).

The python code for the pipeline estimating the π, the π_NS_/π_S_ for all positions and the Tajima’s D of each gene in PoolSeq dataset is available in Github (https://github.com/leveveaudrey/POOLSEQ-analysis-of-polymorphism; https://doi.org/10.5281/zenodo.19051966).

## Acknowledgements

This work has been supported by the Czech Science Foundation (GACR 22-20240S granted to C.L-P). Computational resources were supplied by the project ‘e-Infrastruktura CZ’ (e-INFRA CZ LM2018140) supported by the Ministry of Education, Youth and Sports of the Czech Republic. A.L thanks personally Stella Huynh for our beneficial suggestions and commentaries.

## Author contributions

O.I developed and performed the transcriptional analysis. J-Y.D and A.L developed and performed the coevolution analysis. A.L analysed and interpreted the data. A.L wrote the manuscript. All authors edited the manuscript.

---

**Figure S1:**
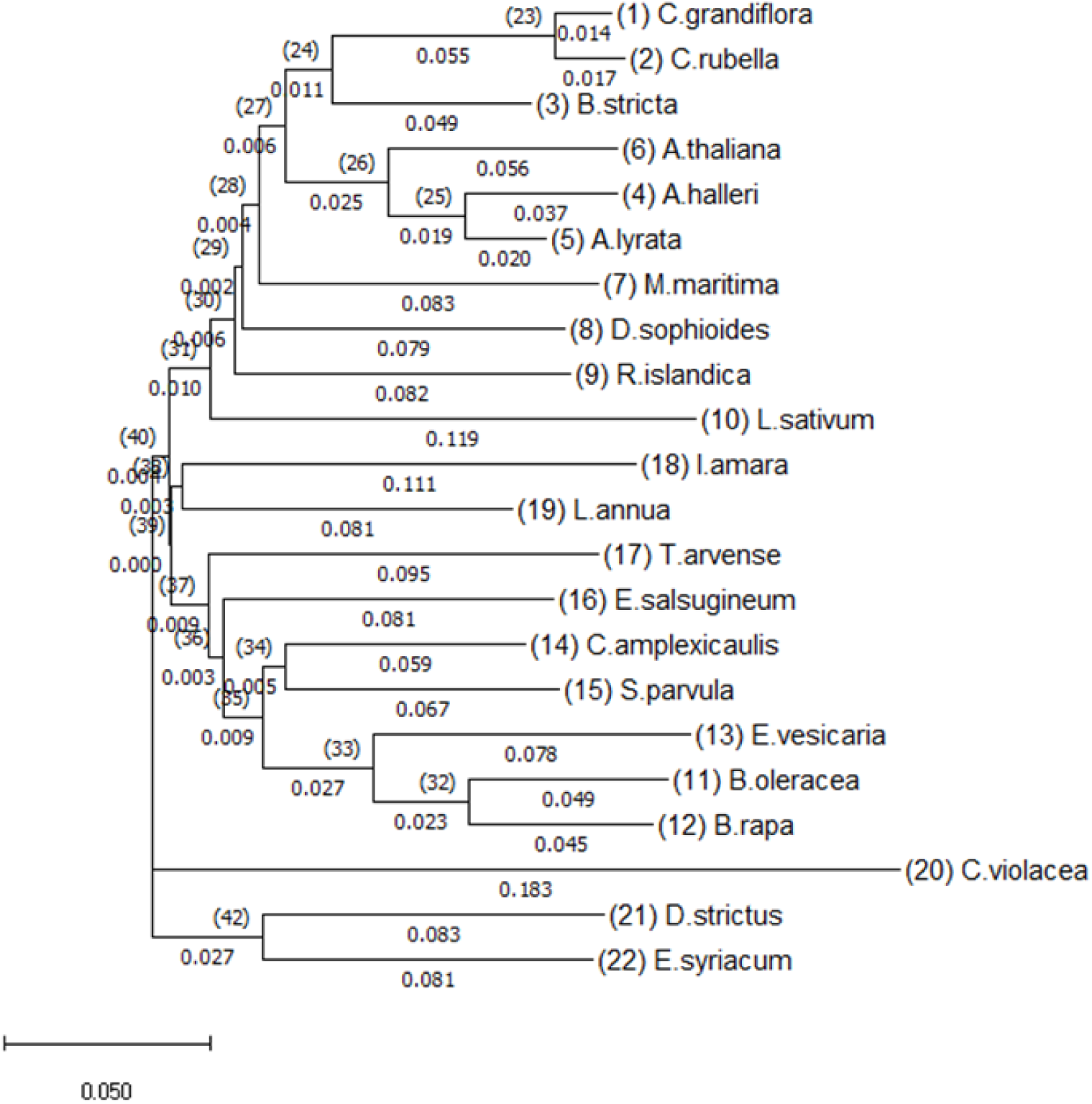
phylogenetic tree obtained by HKY85 model based on the 1055 endosperm genes found in all species after 1000 bootstrap. The percentage of trees in which the associated haplotypes clustered together is shown next to the branches. The tree is drawn to scale, with branch lengths measured in mean number of substitutions per site.

